## Supplemental Data for "Aberrant cortical development is driven by impaired cell cycle and translational control in a *DDX3X* syndrome model"

### Figure S1

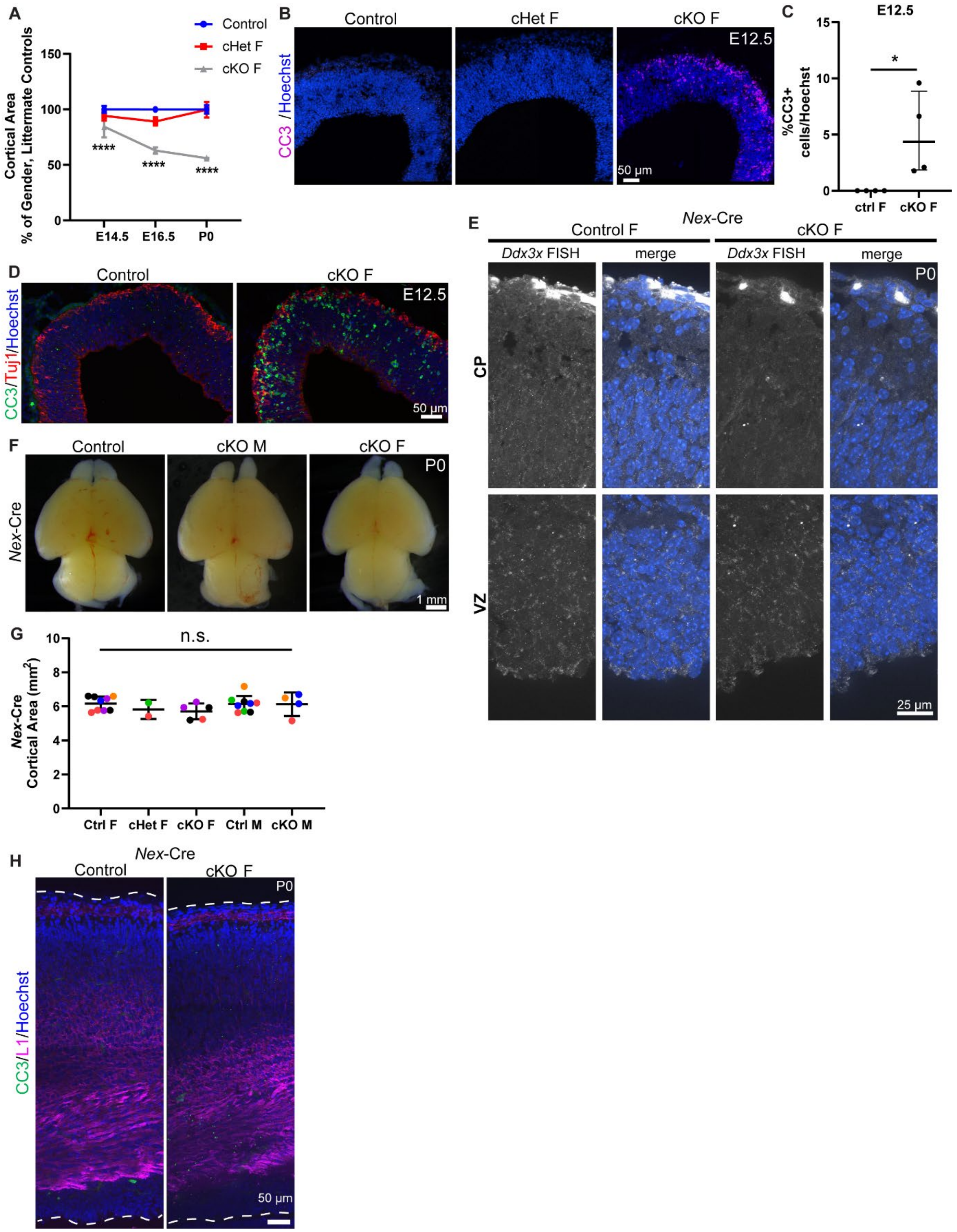

A
Control
cHet F
cKO F
Cortical Area
% of Gender, Littermate Controls
150
100
50
0
E14.5
E16.5
P0
****
B
Control
cHet F
cKO F
E12.5
CC3 /Hoechst
50 µm
C
E12.5
%CC3+ cells/Hoechst
15
10
5
0
ctrl F
cKO F
*
D
Control
cKO F
E12.5
CC3/Tuj1/Hoechst
50 µm
E
Nex-Cre
Control F
cKO F
Ddx3x FISH
merge
P0
CP
VZ
25 µm
F
Control
cKO M
cKO F
P0
Nex-Cre
1 mm
G
n.s.
Nex-Cre Cortical Area (mm²)
10
8
6
4
2
0
Ctrl F
cHet F
cKO F
Ctrl M
cKO M
H
Nex-Cre
Control
cKO F
P0
CC3/L1/Hoechst
50 µm

**Figure S1. *Ddx3x* loss from neural progenitors, but not neurons leads to microcephaly and apoptosis.** (*A*) Temporal quantification of cortical area in control, cHet F, and cKO F at E14.5, E16.5 and P0. (*B*) Representative coronal sections from control, cHet F, and cKO F at E12.5 immunostained with CC3 (magenta) and Hoechst. (*C*) Quantification of CC3+ cells in control and cKO F at E12.5. (*D*) Representative coronal sections from control and cKO F at E12.5 immunostained with CC3 (green), Tuj1 (red) and Hoechst. (*E*) Representative coronal sections of smiFISH for *Ddx3x* mRNA in *Nex*-Cre control and cKO female cortices at P0 showing the ventricular zone (VZ) and cortical plate (CP). (*F*) Representative whole mount images of *Nex*-Cre control and cKO male and female brains at P0. (*G*) Quantification of *Nex*-Cre cortical area at P0. (*H*) Representative coronal sections from *Nex*-Cre control and cKO F at P0 immunostained with CC3 (green), L1 (magenta) and Hoechst.

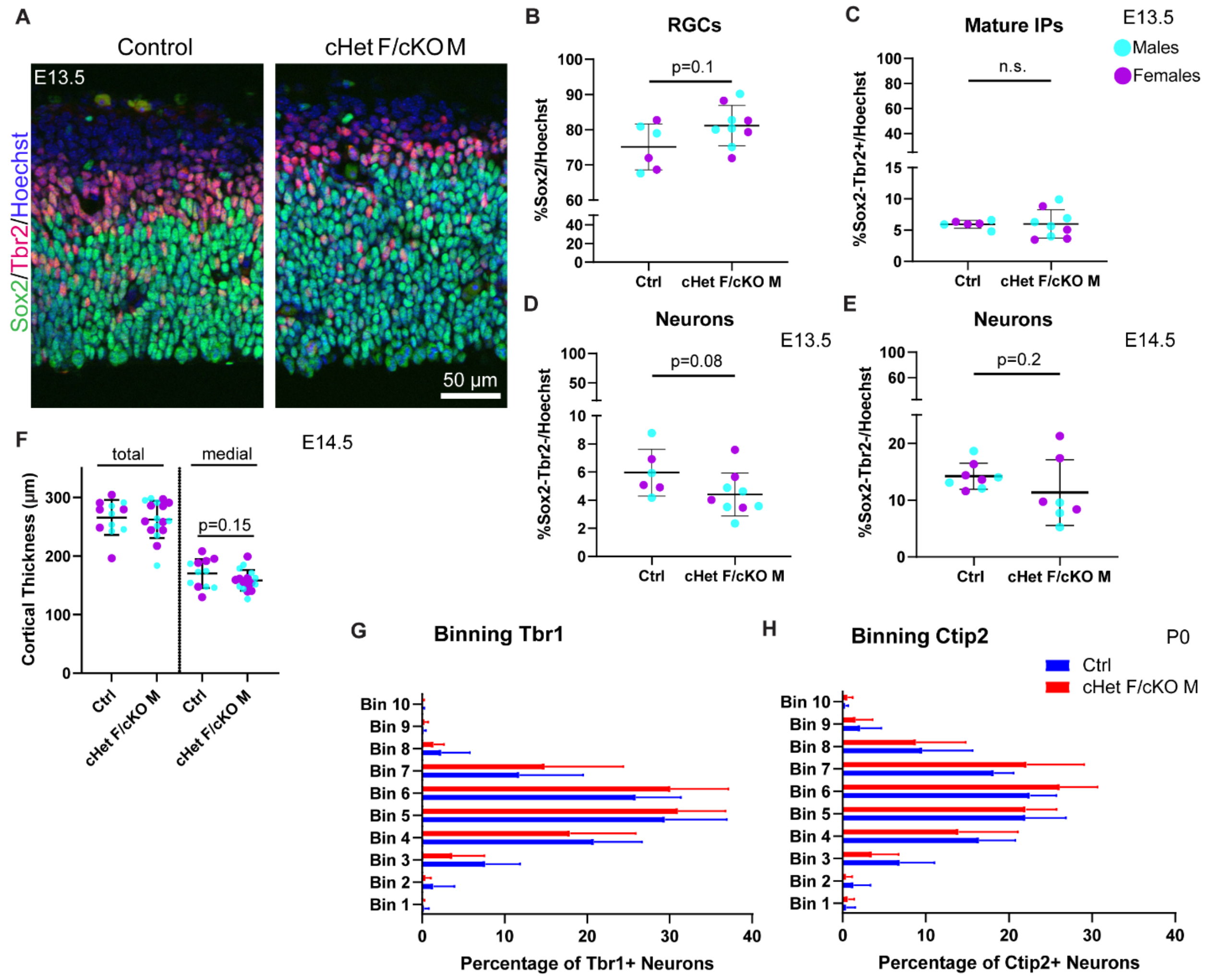


**Figure S2.** ***Ddx3x*** **depletion leads to more progenitors and less neurons at E13.5, but does not affect cortical thickness or laminar position of neurons.** (*A*) Representative coronal sections from control and cHet F/cKO M at E13.5 immunostained with Sox2 (green), Tbr2 (red) and Hoechst (control M and cKO M shown). (*B-D*) Quantification of RGCs (Sox2+) (B), mature IPs (Sox2-,Tbr2+) (*C*), Sox2-Tbr2- ("neurons", *D*) density relative to Hoechst in control and cHet F/cKO M at E13.5. (*E*) Sox2-Tbr2- ("neurons") density relative to Hoechst in control and cHet F/cKO M at E14.5. (*F*) Quantification of cortical thickness at E14.5 in control and cHet F/cKO M measured medially and laterally (total) or just medially. (*G, H*) Quantification of the distribution of Tbr1+ (*G*) and Ctip2+ (*H*) cells in control and cHet F/cKO M at P0.

**Figure S3**

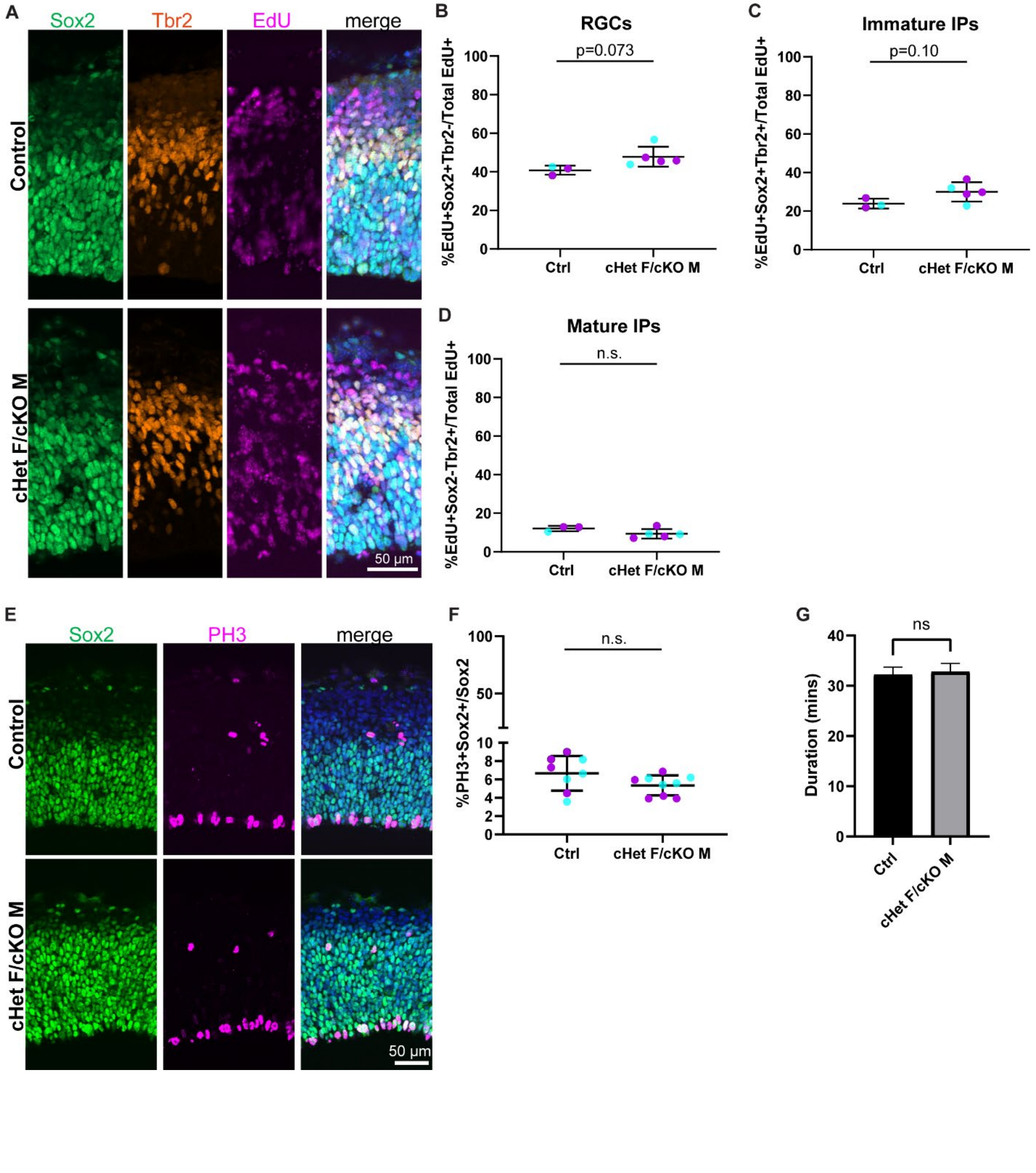
A
Sox2
Tbr2
EdU
merge
Control
cHet F/cKO M
50 µm
B
RGCs
p=0.073
%EdU+Sox2+Tbr2-/Total EdU+
Ctrl
cHet F/cKO M
C
Immature IPs
p=0.10
%EdU+Sox2+Tbr2+/Total EdU+
Ctrl
cHet F/cKO M
D
Mature IPs
n.s.
%EdU+Sox2-Tbr2+/Total EdU+
Ctrl
cHet F/cKO M
E
Sox2
PH3
merge
Control
cHet F/cKO M
50 µm
F
n.s.
%PH3+Sox2+/Sox2
Ctrl
cHet F/cKO M
G
ns
Duration (mins)
Ctrl
cHet F/cKO M

**Figure S3. *Ddx3x* depletion prolongs cell cycle duration in RGCs and immature IPs but does not affect mitosis duration.** (*A*) Representative coronal sections from control and cHet F/cKO M at E14.5 immunostained with Sox2 (green), Tbr2 (red), EdU (magenta) and Hoechst (control F and cHet F shown). (*B-D*) Quantification of RGCs (EdU+Sox2+Tbr2-) (*B*), Immature IPs (EdU+Sox2+Tbr2+) (*C*), Mature IPs (EdU+Sox2-Tbr2+) (*D*), density relative to total EdU+ in control and cHet F/cKO M at E14.5. (*E*) Representative coronal sections from control and cHet F/cKO M at E14.5 immunostained with Sox2 (green), PH3 (magenta), and Hoechst (control M and cKO M shown). (*F*) Quantification of PH3+Sox2+/Sox2+ in control and cHet F/cKO M. (*G*) Quantification of mitosis duration in control and cHet F/cKO M at E14.5 from live imaging analysis.

Figure S4

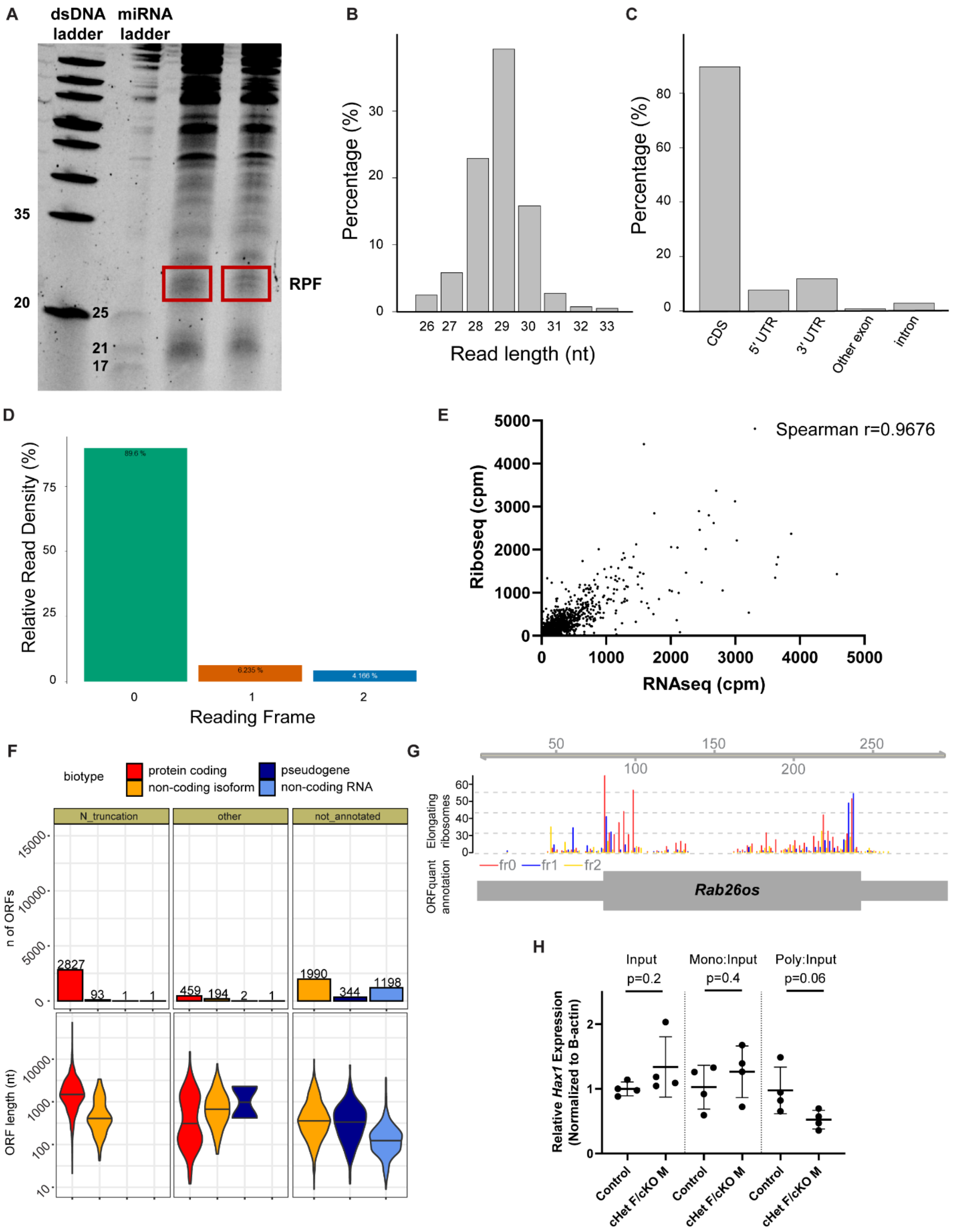
A
dsDNA ladder
miRNA ladder
35
20
25
21
17
RPF
B
Percentage (%)
Read length (nt)
C
Percentage (%)
CDS
5' UTR
3' UTR
Other exon
intron
D
Relative Read Density (%)
Reading Frame
E
Spearman r=0.9676
Riboseq (cpm)
RNAseq (cpm)
F
biotype
protein coding
non-coding isoform
pseudogene
non-coding RNA
N_truncation
other
not_annotated
n of ORFs
ORF length (nt)
2827
93
459
194
1990
344
1198
G
Elongating ribosomes
ORFquant annotation
fr0
fr1
fr2
Rab26os
H
Input
p=0.2
Mono:Input
p=0.4
Poly:Input
p=0.06
Relative Hax1 Expression (Normalized to B-actin)
Control
cHet F/cKO M

**Figure S4. Quality Control Assessment of Ribosome Profiling in *Ddx3x* cKO mice.** (*A*) Representative denaturing urea gel of embryonic cortices treated with RNase I illustrating RPFs (red box). (*B-D*) RibosomeProfilingQC assessment of deep sequencing of cDNA libraries showing read length distribution (B), percent of reads mapping to CDS and UTRs, etc (C), and reading frame (D). (*E*) Comparison of RNAseq cpm and Riboseq cpm using all reads from all transcripts from WT data (excluding non-polyA histone and multi-mapping ribosomal genes); Spearman r=0.9676. 5 transcripts were omitted when reducing axes for readability. (*F*) *De novo* identification of translated ORFs, including Number of detected ORFs with their length (in nucleotides) for different ORF categories and annotated biotypes. (*G*) A novel translated ORF in the lncRNA Rab26os showing the P-sites position colored by frame (middle), and ORFquant-derived annotation (bottom). (*H*) RT-qPCR quantification of mRNA levels for putative Ribo-seq candidate, *Hax1*, in input sample, monosome, and polysome fractions.
